# Consistency Analyses of Open-source Software for Motor Unit Decomposition Using High-density Electromyography Signal

**DOI:** 10.64898/2026.07.07.737019

**Authors:** Jirui Fu, Shiyu Zhang, Helen J. Huang, Mohsen Rakhshan, Yue Wen

## Abstract

Motor unit (MU) decomposition using high-density surface electromyography (HD-sEMG) has been widely used to characterize MU behavior in neurophysiology and to build neural-machine interfaces for wearable robots. Recently, many open-source software tools for MU decomposition have been made available on GitHub, which could reduce the effort of researchers in the field. However, the consistency among these open-source tools has never been studied, making researchers hesitate to use them. In this study, we collected 7 open-source software tools on GitHub and applied them to decompose MUs from an open-source HD-sEMG dataset (including 11 isometric contraction trials) to investigate the consistency among these tools. To create a comprehensive MU pool for reference, we combined all unique MUs identified by seven tools, visually inspected and removed bad MUs, and manually edited all remaining MU spike trains. Across 7 tools for 11 trials, the number of identified MUs ranges from 167 to 736. The number of valid MUs after expert inspection ranges from 29 to 210, which is 10% to 72% of the reference pool. The rate of agreement between the raw MUSTs and the manually edited MUSTs ranges from 0.86 to 0.94, and the averaged number of edits per MU to correct misalignments ranges from 14 to 39. The results show inconsistency in the implementation and procedures of each tool, which results in an inconsistent number of identified MUs and valid MUs (29 vs 210). In general, a substantial amount of effort is required to process the raw MUSTs from each tool to conduct further research analysis. This study provided a guideline for using open-source software tools for MU decomposition and indicated that it would be beneficial to develop tools to automatically edit the MUSTs.

## I. Introduction

The electromyography signal (EMG) represents the superposition of MU action potentials (MUAPs) from each activated MU within the range detectable by electrodes [1]. The motor unit (MU) is the smallest functional unit in the neuromuscular system that contributes to the generation of the EMG signal. They can be decomposed from the high-density surface electromyography signal (HD-sEMG), a variant of the EMG signal recorded using multiple (*>* 4) closely-spaced, noninvasive electrodes [2]. The decomposition of MUs enables the analysis of human neuromuscular system behavior, which has applications in the control of prosthetic devices and the diagnosis of neuromuscular disorders [3].

Currently, blind source separation (BSS) methods for decomposing MUs encompass the convolution kernel compensation algorithms (CKC), which facilitate decomposition by compensating for unknown mixing channels (convolution kernels) [4]. Another method is the Fast Independent Component Analysis (FastICA) algorithm which aims to discover orthogonal rotations of pre-whitened data that optimize a measure of non-Gaussianity [5]. Additionally, an algorithm proposed by Negro et al. [6] integrates the principles of FastICA and CKC to achieve MU decomposition.

So far, numerous software applications have been developed using programming languages such as MATLAB© and Python™ for decomposing MUs. An example is the “DEMUSE Tool” created by Holobar et al. [4], which decomposes MUs using the CKC algorithm; however, this tool is proprietary and needs to be purchased. Conversely, open-source software (OSS) such as “EMG2MU” by Shirazi et al. [7], which utilizes the FastICA [5] with a “peel-off” [8] algorithm, and “EMGdecomPy” by King et al. [9], based on the algorithm described in [6], have been developed for the same purpose. The accuracy of these open-source software solutions has been validated by their developers during the development process. In comparison to proprietary software, the availability of open-source options offers researchers economical alternatives for conducting projects involving MU decomposition, without incurring the costs of proprietary software or necessitating the development of their own programs from scratch.

While open-source software offers advantages in decomposing MUs, several limitations persist. Although these tools have been shown to be capable of identifying MUs, there is a lack of studies benchmarking the consistency across these open-source software using identical datasets and hyperparameters. Furthermore, the MUs identified by BSS methods potentially comprise duplicate MUs or noisy MUs that require removal [3]. Therefore, further inspection grounded in neurophysiological principles is essential to confirm that the identified MUs are indeed valid. However, none of these open-source software has been validated in terms of their ability to identify valid MUs that align with neurophysiological knowledge.

To address these limitations, this paper presents consistency analyses of open-source software designed for MU decomposition using the same dataset and parameters. Furthermore, a rule-based framework [3] is implemented to automatically evaluate the identified MUs and select the valid ones. This study examines the discrepancies in valid MUs identified by different open-source software. The findings are intended to support the selection of open-source software for research in MU decomposition and MU identification.

The organization of the manuscript is as follows: Section II describes the protocol for scanning open-source software and provides a concise description of the methods in this study. The results of the consistency analyses are presented in Section III. A discussion about the causes underlying differences among the selected open-source software and guidelines for choosing and evaluating open-source software for MU decomposition and MU identification is provided in Section IV. Finally, a conclusion is drawn in Section V.

## II. Methods

### A. Description of Dataset

We utilized the open-source dataset from the study by Hug et al. [10] that includes HD-sEMG signals recorded from three muscles: the gastrocnemius lateralis (GL), gastrocnemius medialis (GM), and tibialis anterior (TA), across various contraction intensities. In this dataset, the HD-sEMG signals for the GL and GM muscles were collected during isometric trapezoidal contractions at intensities corresponding to 10%, 30%, and 50% of their maximum voluntary contraction (MVC). However, the data for the TA muscle were recorded with intensities of 35%, 50%, and 70% MVC.

### B. Open-source Software for MU Decomposition

The open-source software incorporated in this study was identified via a search conducted on GitHub using the specified keywords: **“HD-sEMG decomposition”, “HD-EMG decomposition”, “motor unit”, and “sEMG decomposition”**.

The search yielded 86 repositories which were subsequently scanned following the pipeline. After scanning the searched software following this procedure, we ultimately found 7 open-source software being selected for evaluation in this work.

1. Read the description or “readme” file of all found repositories, and excluded 77 repositories that do not have the functionality to decompose MUs.
2. Read through the code of all 9 repositories [7], [9], [11]– [17] to check if any of the repositories use code from other developers and make sure all the repositories are original and complete. From this step, we excluded 2 repositories [16], [17].
3. Modify the input and output of the 7 included repositories so that they are compatible with the open-source testing dataset and return the same format output.

Table I provides a summary of the included open-source software, including a unique ID (the initials of the author and programming language used) for use throughout the manuscript, detailed name of the repositories, the algorithms employed, as well as the programming languages utilized. These software solutions have been developed by various contributors to facilitate MU decomposition from HD-sEMG signals.

**TABLE I.**
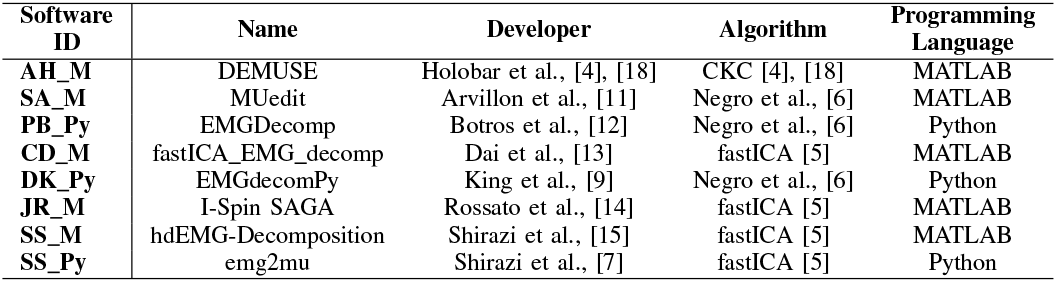
Summary of the included software.

### C. Algorithm for MU Decomposition

Within the framework of MU decomposition algorithm, the raw HD-sEMG signal is denoted as ***X*** *∈* ℝ^*m×N*^, where *m* represents the number of channels and *N* represents the total number of samples. Before running the decomposition algorithm, the raw HD-sEMG signal was preprocessed. First, the raw signal is centered by subtracting its mean value. Then, the centered signal is extended with an extension factor *R*; the resulting centered and extended HD-sEMG signal was denoted as ***Z*** *∈* ℝ^*m*(*R*)*×N*^ . Finally, we whiten the centered and extended HD-sEMG signal ***Z*** using zero-phase component analysis (ZCA), which involves decomposing the covariance matrix Cov(***Z***).

Algorithm 1 stands for the general framework for MU decomposition. We unified the hyperparameter when decomposed the MUs using different software to preserve the consistency, these hyperparameter are shown in Table II. Common terms and notation used in the algorithm were given in Table III. The primary difference between the fastICA algorithm [8] and the one used by Negro et al. [6] was the CKC refinement process. Inside the CKC refinement algorithm, the source signal ***s*** (or IPTs) was calculated by projecting the extended signal on the separation vector (*w*). The source represented the continuous signals whose negative entropy has been maximized. Peak-finding algorithms and K-means algorithms were then applied to the IPTs to extract the discharge time index.

**TABLE II.** Unified Hyperparameter Settings in EMG Decomposition algorithm.

| Notation | Name | Note |
| --- | --- | --- |
| M | Number of Iteration | 150 |
| R or nbextchan | Extension Factor<br>Extended Channle Number | 16<br>1000 |
| $\varepsilon$ | tolerance | $10^{-4}$ |
| SIL_Threshold | SIL_Threshold | 0.6 |

**TABLE III.** Common Terms used in EMG Decomposition Algorithm.

| No. | Notation | Name | Note |
| --- | --- | --- | --- |
| 1 | $w$ | Separation vector | Updated at each iteration |
| 2 | $B$ | Separation matrix | Store separation vector $w$ |
| 3 | IPTs or $\hat{s}$ | Estimated source | Acquired by $w^T Z$ |
| 4 | PT | Discharge time index | Acquired from IPTs |
| 5 | ISI | Inter-spike interval | Acquired from PT |
| 6 | CoV | Coefficient of variation | Acquired from ISI |
| 7 | SIL | Silhouette score | Use to evaluate performance in 3 |
| 8 | PNR | Pulse-noise ratio | Use to evaluate performance in 3 |

### D. Evaluation Protocols

We employed the included open-source software to decompose MUs from the open-source dataset presented in [10]. The decomposed MU were counted and referred to as “Identified” shown in Table IV. Next, we combined the unique MUs from all software and performed visual inspection and editing using DEMUSE tool software (v4.9; The University of Maribor, Slovenia) developed on MATLAB [18]. The edited MUs served as a reference for evaluation.

**TABLE IV.** Total Number of MU after each stage of Operation.

| Software | Number of MUs |  |  |  |  |  |
| --- | --- | --- | --- | --- | --- | --- |
|  | Identified | Repeated | Noisy | Potential | Valid | Ratio |
| AH_M | 130 | 0 | 0 | 130 | 130 | 45% |
| SA_M | 292 | 31 | 4 | 257 | 188 | 64% |
| PB_Py | 167 | 38 | 89 | 40 | 29 | 10% |
| CD_M | 226 | 42 | 23 | 161 | 142 | 49% |
| DK_Py | 736 | 493 | 189 | 54 | 50 | 17% |
| JR_M | 355 | 40 | 41 | 274 | 210 | 72% |
| SS_M | 196 | 15 | 20 | 161 | 141 | 48% |
| SS_Py | 505 | 331 | 17 | 157 | 137 | 47% |
| Manual |  |  |  | 377 | 292 |  |

#### Algorithm 1

Algorithm for MU Decomposition [6]

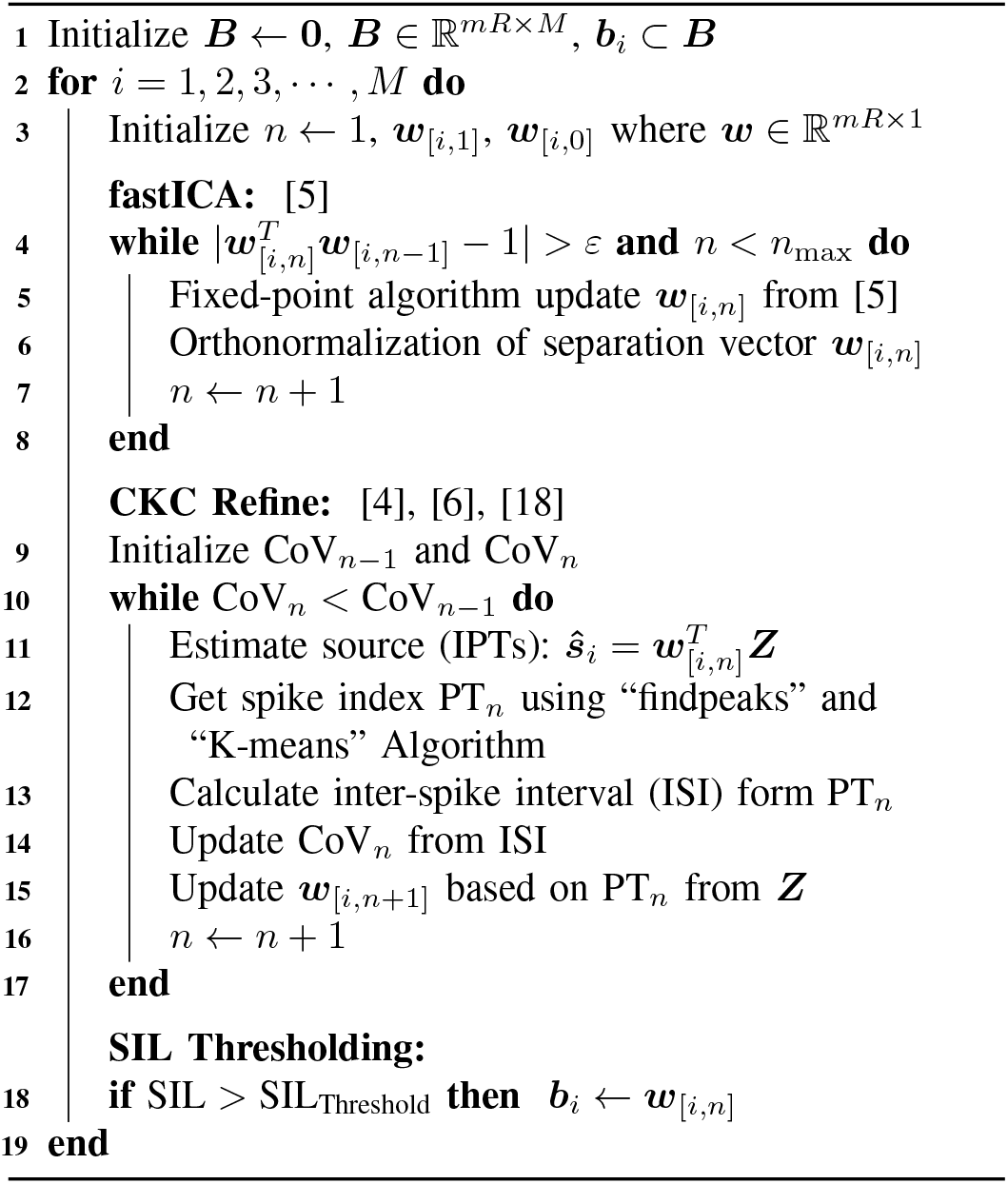

Given that the raw MUs decomposed by the included software were susceptible to duplication due to the extension of the HD-sEMG signal and noise associated with artifacts, we performed a post-processing stage to eliminate both duplicate and noisy MUs. To address the issue of duplicate MUs and spikes, we adhered to the criteria outlined in [19], which can be succinctly described as:

- If any pair of MU spike trains (MUST) are triggered within *±*1 ms, this pair of MUST would be considered as synchronized.
- If more than 30% of MUSTs of two MUs are synchronized, this pair of MUs would be considered as duplicated, and one of them needs to be removed.

Subsequently, we removed the synchronized/duplicated MUs, and updated the corresponding pulses in the IPTs. The removed MUs were counted and referred to as “Repeated” shown in Table IV. Then, the processed MUs were analyzed to identify and exclude any noisy MUs. For the removal of noisy MUs, we re-computed the pulse-noise-ratio (PNR) and Silhouette score (SIL) and removed MUs with PNR*<*25 and SIL*<*0.6. We used a low PNR threshold and a low SIL threshold in this work to keep more potential MUs for further manual inspection and editing. The removed MUs were counted and referred to as “Noisy” shown in Table IV. The MUs left after removing “Repeated” and “Noisy” were counted and referred to as “Potential” shown in Table IV.

Following the post-processing of raw MUs, we extracted the unique MUs from all software to generate a comprehensive MU pool (including 377 MUs), and we further visually inspected and manually edited all potential unique MUs in the MU pool. After manual editing and refinement, MUs with PNR lower than 28 will be removed from the MU pool, resulting in 292 MUs. Mapping the MU pool back to the MUs identified by each software, the preserved MUs from each software after operator manual editing were counted and referred to as “Valid” shown in Table IV. We also calculated the ratio of the number of “Valid” MUs from each software to that of the unique MUs pool (i.e., 292 MUs) as shown in the “Ratio” column in Table IV. The outcomes of this benchmark served as an evaluation of the quality of the open-source software in MU decomposition. We also compared the raw MUSTs from each software with the manually edited MUSTs to check the performance of each software. Specifically, we calculated the number of false positives (FPs) and false negatives (FNs) and the rate of agreement (RoA; details in the following section) between the paired raw MUST and the edited MUST.

Finally, we assessed the consistency of the software by computing the RoA for each MU between spike trains produced by any two software, namely software A and software B, utilizing the following equation:

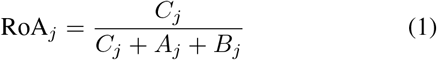

where *C*_*j*_ denotes the number of common spikes of the *j*th MU decomposed by both software that were triggered within *±*1*ms* window [6], *A*_*j*_ is the number of spikes that were decomposed by software A but not by software B, and *B*_*j*_ is the number of spikes that were decomposed by software B but not by software A. We used the same method for detecting duplicate MUs [19] with a threshold of 50% to identify common MUs between any two software tools.

## III. Results

### A. Presence of Duplicate and Noisy MUs in Decomposed MUs by Open-source Software

Table IV lists the number of MUs identified by each open-source software, as well as the number of duplicate and noisy MUs following post-processing. The “Potential” column indicates the number of MUs after removing duplicate and noisy MUs, and the unique MUs across all software are 377 for further visual inspection and manual editing by an expert. The “Valid” column reflects the number of remaining MUs of each software tool after expert inspection and removing bad MUs with PNR lower than 28, and thus the number of unique MUs is reduced from 377 to 292. The “Ratio” column calculates the ratio of the number of “Valid” MUs from each software to that in the MU pool (i.e., 292 MUs). A ratio lower than 45% (as of “AH M”) was marked as red. According to the data presented in Table IV, a significant discrepancy exists between the number of decomposed MUs from each software and the number of valid MUs after expert inspection. The findings reveal that “DK Py” by King et al. decomposed the largest number of MUs (736 MUs). However, it only revealed 17% of the unique MU pool, and a large number of MUs were discarded (682 MUs) due to duplication or noise. “PB Py” by Botros et al. got a small number of identified MUs (167 MUs) and revealed only 29 valid MUs (10% of the unique MU pool). We also observed “JR M” by Rossato et al. produced the highest number of valid MUs (210 MUs, 72% of the unique MU pool) and the software “SA M” by Arvillon et al. also yields a higher number of valid MUs (188 MUs, 64% of the unique MU pool).

### B. Substantial Quantity of Manual Edits is Required for MUs Decomposed by Open-source Software

Figure 1 shows that the average RoA between the raw MUSTs from each software tool (Figure 1(A)) and the manually edited MUSTs ranges from 0.86 to 0.94 and that the averaged number of edits per MU to correct FPs and FNs ranges from 14 to 39 (Figure 1(B)) . Even though a large number of MUs were discarded, “DK Py” by King et al. prevails over other software by having the highest RoA (0.94) and the least number of edits (14 edits). “SA M” by Arvillon et al. and “JR M” by Rossato et al. have a low RoA (0.86 and 0.86, respectively) and a high number of edits (39 edits and 37 edits, respectively).

**Fig. 1.**
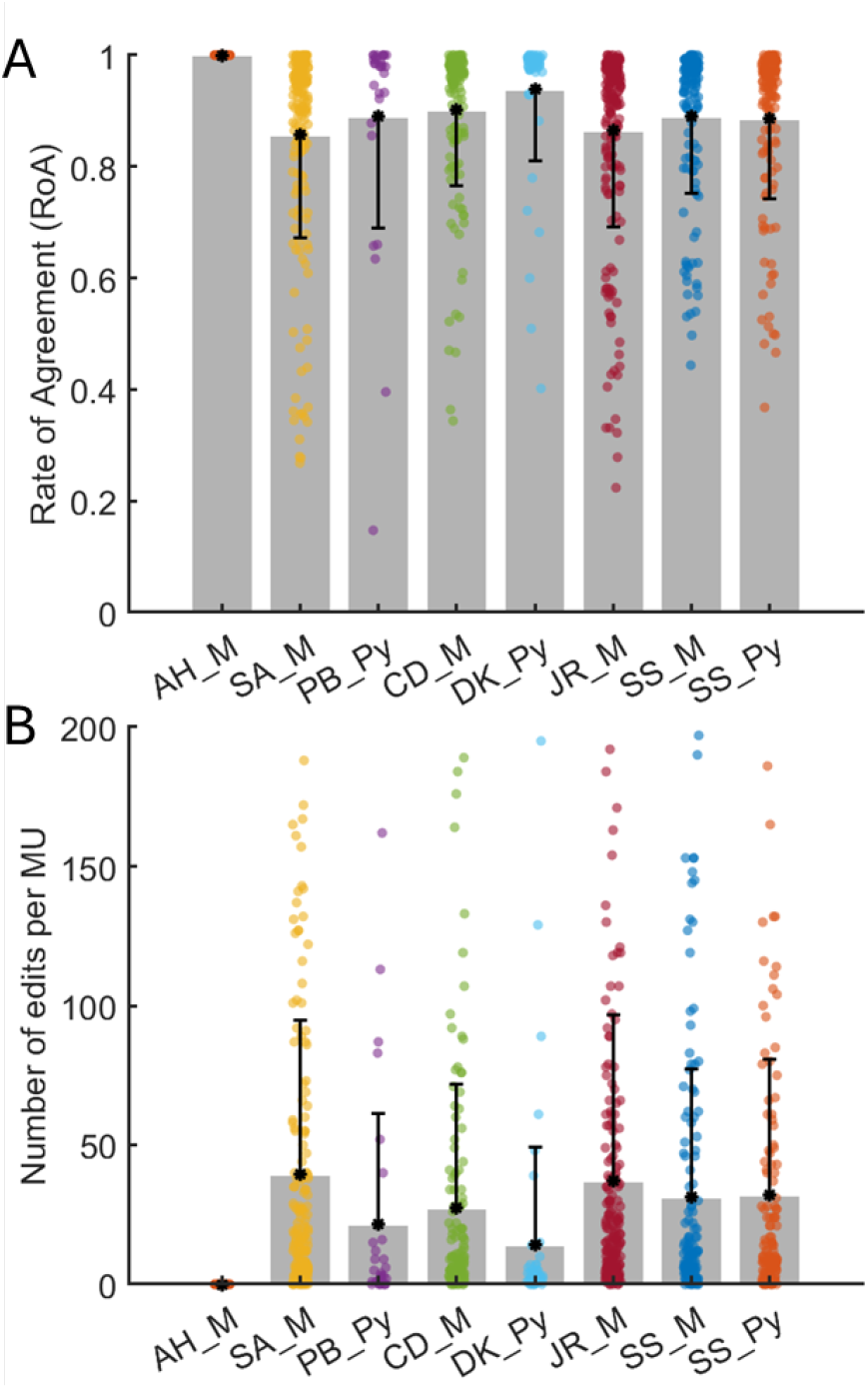
A: Rate of agreement (RoA) between the raw MUSTs from each tool and the manually-edited MUSTs, with 1 indicating a perfect match. Each dot indicates one MU, and the number of dots (MUs) is reported as “Valid” in Table IV. The error bar is calculated using all dots (MUs). B: The number of edits per MU to refine the raw MUSTs from each tool to match the manually-edited MUSTs, with 0 indicating a perfect match. The error bar is calculated using all dots (MUs).

### C. Consistency Among the Open-source Software in MU Decomposition

Table V shows the RoA between each pair of software calculated based on the number of overlapped MUs, the averaged RoA across these software is 0.91, indicating inconsistency in the MUSTs decomposed by these software. The software “DK Py”, developed by King et al. [9] had the lowest number of common MUs with other software (23 MUs) and exhibited the highest RoA (0.94) when compared with other software, indicating it only kept the high-quality MUs that require minimal editing. Conversely, the software “JR M”, by Rossato et al. [14] had the highest number of common MUs with other software (103 MUs) but generated the lowest RoA (0.88), as it includes 210 MUs with some low-quality MUs requiring manual editing.

**TABLE V.** Rate of Agreement (ROA) and Number of Overlapping MUS between each pair of algorithms.

|  | AH_M | SA_M | PB_Py | CD_M | DK_Py | JR_M | SS_M | SS_Py |
| --- | --- | --- | --- | --- | --- | --- | --- | --- |
| AH_M | 130 | 107 | 26 | 92 | 45 | 105 | 96 | 87 |
| SA_M | 0.93 | 188 | 25 | 129 | 44 | 161 | 124 | 120 |
| PB_Py | 0.94 | 0.91 | 29 | 22 | 20 | 24 | 23 | 23 |
| CD_M | 0.93 | 0.85 | 0.88 | 142 | 42 | 128 | 125 | 116 |
| DK_Py | 0.95 | 0.96 | 0.95 | 0.97 | 50 | 44 | 42 | 40 |
| JR_M | 0.93 | 0.85 | 0.90 | 0.87 | 0.90 | 210 | 130 | 126 |
| SS_M | 0.92 | 0.89 | 0.88 | 0.90 | 0.95 | 0.87 | 141 | 116 |
| SS_Py | 0.92 | 0.86 | 0.87 | 0.90 | 0.95 | 0.85 | 0.90 | 137 |

## VI. Discussion

This research examined the consistency of open-source software tools for decomposing MUs, using an identical open-source dataset. The study focused on assessing the number of valid MUs identified, the quantity of manual modifications required, and the rate of agreement between MUSTs decomposed by two open-source software packages. Additionally, a comparative analysis was conducted between the quality and consistency of these open-source software tools and a closed-source reference software, DEMUSE, which applies CKC algorithms.

To achieve high-quality MUs, the units decomposed by each open-source software required substantial post-processing to remove duplicated and noisy MUs, as well as to edit false negatives and false positives. As indicated in Tables IV and V in Section III, between 18% and 90% MUs were proved valid within each software’s output; furthermore, each valid MU produced by the open-source tools required from 20 to 50 manual edits on average to correspond to the MU manually edited by an expert. In contrast, the DEMUSE software includes an approach to remove duplicate and noisy motor units which have been widely accepted by other researchers [19]. Upon investigating the cause of this discrepancy, it was discovered that the reference software comprises a module for removing duplicates following the pipeline introduced in [19], capable of eliminating duplicate MUs if the number of synchronized MU spike trains (MUSTs) exceeds 50% of the total MUSTs, and modifies the retained MUs to eliminate synchronized MUSTs. Conversely, none of the evaluated open-source software is equipped and used with a comparable function to remove duplicates and edit synchronized MUSTs, resulting in considerable numbers of duplicate MUs being detected during post-processing. Furthermore, regarding the presence of noisy MUs, it is hypothesized that this issue arises from imperfect whitening of the input HD-sEMG signal, which is the initial step in all BSS algorithms. However, further investigation will be necessary to explicate the source of the noise. Finally, the reason for the discrepancy in the number of manual edits required by open-source software and reference software remains unidentified due to the proprietary nature of the reference, rendering the detailed post-processing protocols of the decomposed MUs inaccessible.

Moreover, the findings revealed inconsistency among the included open-source software, even those employing the same algorithm and programming language. For instance, the RoAs between pair CD M-JR M and CD M-SS M are about 88%. Similar inconsistencies are also observed between other pairs of open-source software tools. It is suspected that these inconsistencies are attributable to the differences in the implementation of BSS algorithms within programming languages. Notably, in comparison to MATLAB, open-source software developed in Python, such as PB Py, CD M, and SS Py, exhibits higher inconsistency with other software. This phenomenon is believed to stem from the open-source nature of Python. Unlike MATLAB, where mathematical operations are developed, tested, and maintained by MathWorks Company, which guarantees inherent reliability for calculation, Python boasts numerous modules written by various developers that perform the same mathematical operations. Moreover, the round-off error between using “float64”(MATLAB) and “float32”(Python packages) will mainly affect the numerical stability of the decomposition due to the IEEE 754 standard [20], especially in calculating the orthogonality of the separation vectors(i.e. Grand-Schmidt Orthogonalization process) [21]. Thus, it is posited that the numerical precision of Python modules for requisite mathematical operations for BSS might affect the consistency of MU decomposition software developed by different individuals, even under the same algorithm. Nonetheless, this hypothesis requires further investigation for validation.

Additionally, it was possible to identify other repositories such as those by Gibbs et al. and Ricioli et al. However, due to the discontinuities and intricate wrapping of data structures, no output could be obtained from their work, prompting the decision to refrain from reporting these two packages until further development occurs.

## V. Conclusion

Our work quantitatively investigated seven open-source MU decomposition software tools using the same open-source dataset under the same hyperparameters. We investigated three main factors: the number of valid MU, the average number of edits, and RoA between results from any two software tools. With the presented results, we conclude that: 1) The number of MU from MU decomposition software tools varied significantly, ranging from 29 to 210 MUs. 2) The number of edits across the seven software tools ranges from 14 to 40 edits per MU, and it is still a burdensome task for researchers to manually correct these FPs and FNs. 3) The RoA among seven open-source software tools ranges between 80%-90%. 4) With the provided benchmark in this work, researchers could have a basic understanding of the reliability and performance of different software. The future work of this study will focus on identifying the causes of the observed discrepancies in number of motor units decomposed among the included open-source software tools. In addition, we will expand the scope of our investigation by exploring and incorporating more open-source software for motor unit decomposition, as well as employing additional datasets collected from different muscles and participants to further evaluate the performance of these tools. Finally, we aim to provide the research community with practical guidelines for implementing and developing open-source software for motor unit decomposition.

## References

[1] Y. Wen, S. Avrillon, J. C. Hernandez-Pavon, S. J. Kim, F. Hug, and J. L. Pons, “A convolutional neural network to identify motor units from high-density surface electromyography signals in real time,” Journal of Neural Engineering, vol. 18, no. 5, p. 056003, apr 2021.

[2] J. Fu, H. J. Huang, and Y. Wen, “Evaluating convolution neural network architecture for neural drive decoding from high-density surface electromyography,” in 2025 International Conference On Rehabilitation Robotics (ICORR), 2025, pp. 1–6.

[3] A. Del Vecchio, A. Holobar, D. Falla, F. Felici, R. Enoka, and D. Farina, “Tutorial: Analysis of motor unit discharge characteristics from high-density surface emg signals,” Journal of Electromyography and Kinesiology, vol. 53, p. 102426, 2020.

[4] A. Holobar and D. Zazula, “Multichannel blind source separation using convolution kernel compensation,” IEEE Transactions on Signal Processing, vol. 55, no. 9, pp. 4487–4496, 2007.

[5] A. Hyvarinen, “Fast and robust fixed-point algorithms for independent component analysis,” IEEE transactions on Neural Networks, vol. 10, no. 3, pp. 626–634, 1999.

[6] F. Negro, S. Muceli, A. M. Castronovo, A. Holobar, and D. Farina, “Multi-channel intramuscular and surface emg decomposition by convolutive blind source separation,” Journal of neural engineering, vol. 13, no. 2, p. 026027, 2016.

[7] S. Y. Shirazi, “emg2mu: Gpu-accelerated high-density emg decomposition,” 2024. [Online]. Available: https://github.com/neuromechanist/emg2mu

[8] M. Chen and P. Zhou, “A novel framework based on fastica for high density surface emg decomposition,” IEEE Transactions on Neural Systems and Rehabilitation Engineering, vol. 24, no. 1, pp. 117–127, 2015.

[9] D. King, J. Ortega, R. Rudyak, and R. Sivanandam, “Emgdecompy,” 2022. [Online]. Available: https://github.com/The-Motor-Unit/EMGdecomPy

[10] F. Hug, S. Avrillon, A. Del Vecchio, A. Casolo, J. Ibanez, S. Nuccio, J. Rossato, A. Holobar, and D. Farina, “Analysis of motor unit spike trains estimated from high-density surface electromyography is highly reliable across operators,” Journal of Electromyography and Kinesiology, vol. 58, p. 102548, 2021.

[11] S. Avrillon, F. Hug, C. Gibbs, and D. Farina, “Tutorial on muedit: an open-source software for identifying and analysing the discharge timing of motor units from electromyographic signals. biorxiv 548568,” 2023.

[12] P. Botros, “emgdecomp,” 2021. [Online]. Available: https://github.com/carmenalab/emgdecomp/

[13] X. Jiang, X. Liu, J. Fan, X. Ye, C. Dai, E. A. Clancy, M. Akay, and W. Chen, “Open access dataset, toolbox and benchmark processing results of high-density surface electromyogram recordings,” IEEE Transactions on Neural Systems and Rehabilitation Engineering, vol. 29, pp. 1035–1046, 2021.

[14] J. Rossato, F. Hug, K. Tucker, L. Lacourpaille, D. Farina, and S. Avrillon, “I-spin live: An open-source software based on blind-source separation for decoding the activity of spinal alpha motor neurons in real-time,” Elife, vol. 12, 2023.

[15] S. Y. Shirazi, “hdemg-decomposition,” 2022. [Online]. Available: https://github.com/neuromechanist/hdEMG-Decomposition

[16] C. Gibbs, “Emg decomposition,” 2021. [Online]. Available: https://github.com/ciaragibbs/EMGDecomposition

[17] G. Ricioli, “semg-decomposition,” 2021. [Online]. Available: https://github.com/guilhermerc/semg-decomposition

[18] A. Holobar and D. Zazula, “Gradient convolution kernel compensation applied to surface electromyograms,” in International Conference on Independent Component Analysis and Signal Separation. Springer, 2007, pp. 617–624.

[19] A. Holobar, M. A. Minetto, A. Botter, F. Negro, and D. Farina, “Experimental analysis of accuracy in the identification of motor unit spike trains from high-density surface EMG,” IEEE Trans. Neural Syst. Rehabil. Eng., vol. 18, no. 3, pp. 221–229, Jun. 2010.

[20] “Ieee standard for floating-point arithmetic,” IEEE Std 754-2019 (Revision of IEEE 754-2008), pp. 1–84, 2019.

[21] N. J. Higham, Accuracy and stability of numerical algorithms. SIAM, 2002.

